# ZINC-FINGER MOTIFS UNIQUE TO ARABIDOPSIS THALIANA PARALOGS RPA1C AND RPA1E ARE REQUIRED FOR RPA-DEPENDENT DNA REPAIR

**DOI:** 10.64898/2026.01.26.701817

**Authors:** Ian A. Mills, Kevin M. Culligan

## Abstract

RPA is a heterotrimeric ssDNA binding protein that is highly conserved across all eukaryotes. Arabidopsis (*Arabidopsis thaliana*) has five RPA1 paralogs divided into three groups (A, B, C) each with unique functions in DNA replication and repair. The group C paralogs (RPA1C and RPA1E in Arabidopsis) function specifically in DNA-damage repair and carry a C-terminal extension unique to group-C paralogs. This C-terminal extension contains a zinc finger motif (ZFM) that is highly conserved and is therefore predicted to be critical to the functionality of the paralogs during DNA damage repair. To address this, we employed a CRISPR-Cas9 strategy to specifically remove the ZFM from RPA1C or RPA1E while leaving the genes otherwise intact (termed C-ZFKO and E-ZFKO). C-ZFKO and E-ZFKO lines were challenged with DNA damaging agents, and their susceptibility was compared to both WT (Col-0) lines and to previously characterized T-DNA null mutants (*rpa1c* and *rpa1e*). To address the role of the respective ZFMs in homologous recombination pathways (HRR), we employed a GUS-reporter system to compare WT lines to C-ZFKO and E-ZFKO lines. We find here that C-ZFKO and E-ZFKO lines displayed hypersensitivity to DNA damaging agents at a level comparable to previously characterized T-DNA null mutants (*rpa1c* and *rpa1e*). When studying the rate of HRR, both C-ZFKO and *rpa1c* showed a drastic reduction in single-strand annealing (SSA) while E-ZFKO and *rpa1e* had a more modest, but still significant decrease. All mutant lines had a comparable decrease in synthesis-dependent strand annealing (SDSA) compared to WT. Thus, we show here that the respective RPA1C and RPA1E-encoded ZFM is crucial for the ability of each paralog to function during DNA damage repair.

## Introduction

Replication protein A (RPA) is a heterotrimeric single-stranded DNA (ssDNA)-binding protein consisting of three subunits: RPA1 (∼70 kDa), RPA2 (∼32 kDa), and RPA3 (∼14 kDa). A leading role of RPA is to protect ssDNA from unwanted interactions such as degradation and hairpin formation (Pal & Levy, 2019). RPA acts across a wide variety of cellular contexts, including DNA replication, recombination, and DNA damage repair (Wold, 1997). The RPA heterotrimer binds with extremely high (subnanomolar) affinity to ssDNA through the use of six distinct DNA-binding domains (DBDs), designated DBD A-F, which can be found across the three subunits (Kim et al., 1994). The DBDs of RPA are also capable of interacting with numerous protein targets, which is crucial for the ability of RPA to function within multiple pathways that involve ssDNA (Chen & Wold, 2014).

One crucial pathway in which RPA binds is DNA damage repair. Two key factors in the DNA-damage response are the checkpoint kinases ataxia-telangiectasia mutated (ATM) and ataxia-telangiectasia and Rad3-related (ATR) (Zhou & Elledge, 2000) which serve as the primary signal transducers in DNA damage repair (Gimenez & Manzano-Agugliaro, 2017). ATR is a more generalized factor while ATM is specifically involved in the repair of DSBs and is recruited by the Mre11-Rad50-Nbs1 (MRN) complex (Cimprich & Cortez, 2008; Lee & Paull, 2005). Recruitment of ATR does not require the MRN complex but instead involves ATR-interacting protein (ATRIP), as well as RPA (Stracker & Petrini, 2011; Zou & Elledge, 2003). RPA-coated ssDNA serves as a binding target for ATRIP, which then recruits ATR (Ball et al., 2007). Following the binding of either ATR or ATM, a signaling cascade is initiated which causes the cell cycle to pause while the cell prepares to either repair the damage or initiate programmed cell death (Amiard et al., 2011; Q. Liu et al., 2000; Matsuoka et al., 1998; Reinhardt et al., 2007). Initiating programmed cell death is a last resort but is sometimes necessary due to the possible consequences of a DSB (Roy, 2014). DSBs are the most deleterious form of DNA damage and can lead to chromosome loss or truncation. However, not all DSBs result in programmed cell death. Instead, multiple complex DNA repair pathways can be utilized to repair DSBs (West et al., 2004).

After the recognition of a DSB and the initiation of the signaling cascade by ATM and ATR, cellular DSB repair generally involves one of three main mechanisms: non-homologous end joining (NHEJ), microhomology-mediated end joining (MMEJ), or homologous recombination repair (HRR) (Figure 1). Of the three methods, NHEJ is the most commonly used but is error-prone, and can result in the alteration or loss of the DNA sequence (Kirik et al., 2000; Roy, 2014; Salomon & Puchta, 1998). MMEJ was previously considered to be a backup pathway for NHEJ, but has recently been shown to have a larger, more consistent role in DSB repair, and also often results in the loss of DNA sequence (Doonan & Sablowski, 2010; Nussenzweig & Nussenzweig, 2007). HRR, while less commonly used than NHEJ, is still a critical repair mechanism of DSBs and can repair DSBs with no loss of DNA sequence. There are two variants of HRR: single-strand annealing (SSA) and synthesis-dependent strand annealing (SDSA; Puchta, 2004). SSA, like NHEJ and MMEJ, is a non-conservative repair mechanism and can result in the loss of kilobases of DNA (Mendez-Dorantes et al., 2018). This contrasts with SDSA, which is a conservative repair mechanism and can repair DNA damage with no alteration of the DNA sequence. Both mechanisms begin with the initiation of a signaling cascade by ATM and ATR, followed by the resectioning of the ends of the DSB by the MRN complex, leaving 3’ OH overhangs (Mannuss et al., 2010). RPA then binds these regions of single-stranded DNA to protect the DNA from harm and any unwanted interactions, while promoting appropriate DNA repair pathways.

**Figure 1.**
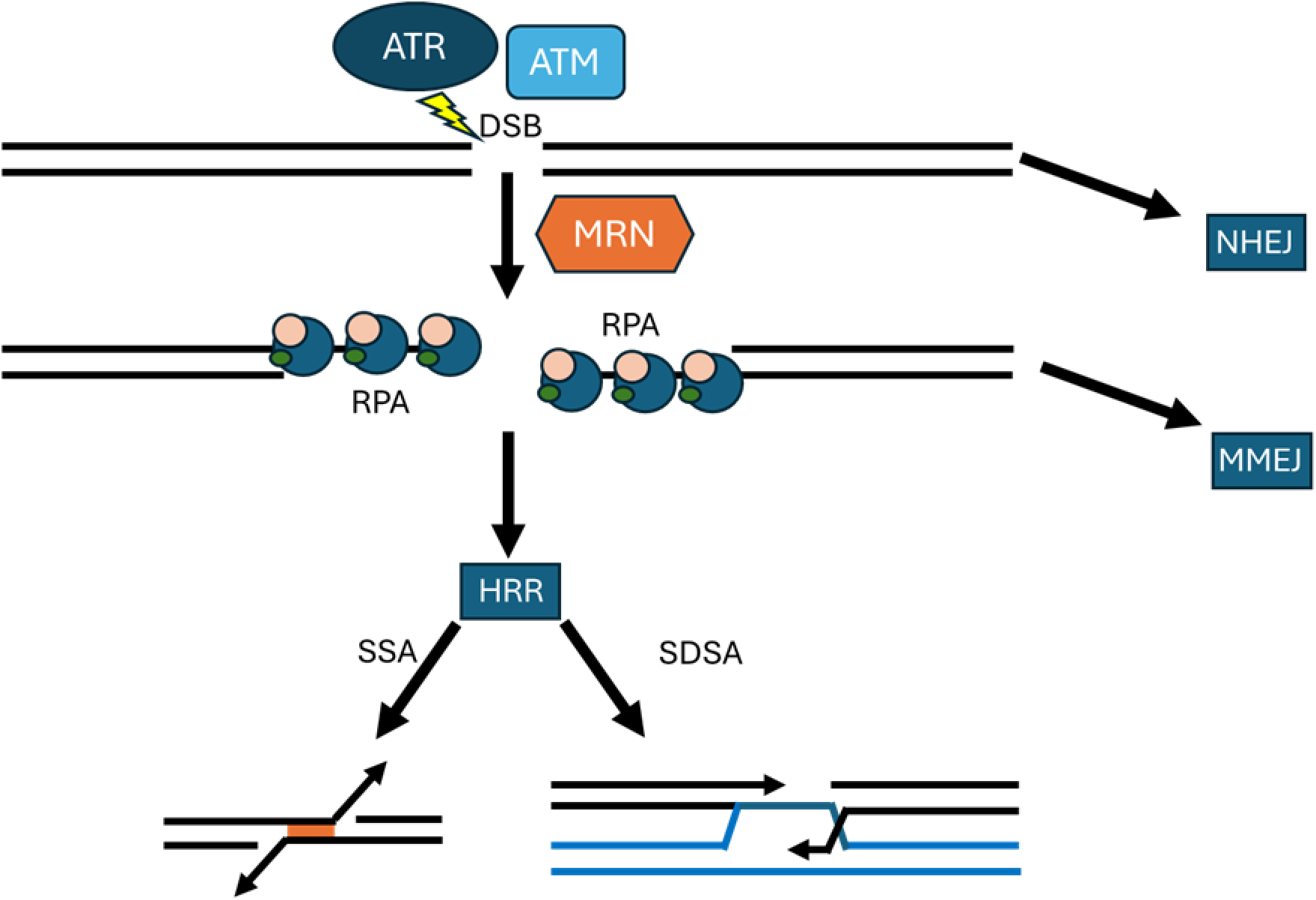
Simplified model of double-strand break repair in plants. Ataxia-Telangiectasia Mutated (ATM) and Ataxia-Telangiectasia and Rad 3 Related (ATR) initiate a signaling cascade. The signaling results in either the direct repair of the break by non-homologous end joining (NHEJ) or the resecting of the ends of the break by the Mre11-Rad50-NBS1 (MRN) complex to create 3’ OH overhangs which are bound by Replication Protein A (RPA). At this point, the cell either repairs the break via microhomology-mediated end joining (MMEJ) or through one of the two versions of homologous recombination repair (HRR): single-strand annealing (SSA) or synthesis dependent strand annealing (SDSA).

With a few exceptions, plants have multiple paralogs of each RPA subunit, while animals and yeast typically have a single version of each subunit. While all plants appear to have multiple RPA subunit paralogs, the number of paralogs of each subunit varies between plants (Shultz et al., 2007). Rice has three paralogs of both RPA1 and RPA2 and only a single RPA3 variant, while *Arabidopsis thaliana* has five RPA1 paralogs (RPA1A, RPA1B, RPA1C, RPA1D, and RPA1E) and two each of RPA2 (RPA2A and RPA2B) and RPA3 (RPA3A and RPA3B) (Aklilu et al., 2014; Ganpudi & Schroeder, 2011; Ishibashi et al., 2006). The Arabidopsis RPA1 paralogs have overlapping, but distinct functionality (Aklilu et al., 2014). The five Arabidopsis RPA1 paralogs fall into one of three groups by function. Group A paralogs (RPA1A) are primarily involved in the progression of meiosis, group B paralogs (RPA1B and D) are primarily involved in replication, and group C paralogs (RPA1C and E) are primarily involved in DNA damage repair.

Arabidopsis RPA1 paralogs are structurally very similar (Aklilu & Culligan, 2016). All five paralogs maintain the same four DNA binding domains (DBD-A, DBD-B, DBD-C, and DBD-F) but the subdomain Binding Surface I (BS-I) can only be found in RPA1A, RPA1C, and RPA1E (Figure 2). BS-I is required for functionality during DNA damage repair but unnecessary for activity during DNA replication (Haring et al., 2008; Longhese et al., 1994; Umezu et al., 1998). The other distinguishing feature between the paralogs is the C-terminal extension, which is found only in group C paralogs and only in plants. This region of ∼176 amino acids in RPA1C paralogs and ∼119 amino acids in RPA1E paralogs always contains at least one CCHC-type (CX2CX4HX4C) zinc finger motif (ZFM) which is hypothesized to be crucial to the functionality of the paralog during DNA damage repair (Aklilu & Culligan, 2016). This region may also be involved in the regulation of DNA damage response in plants.

**Figure 2.**
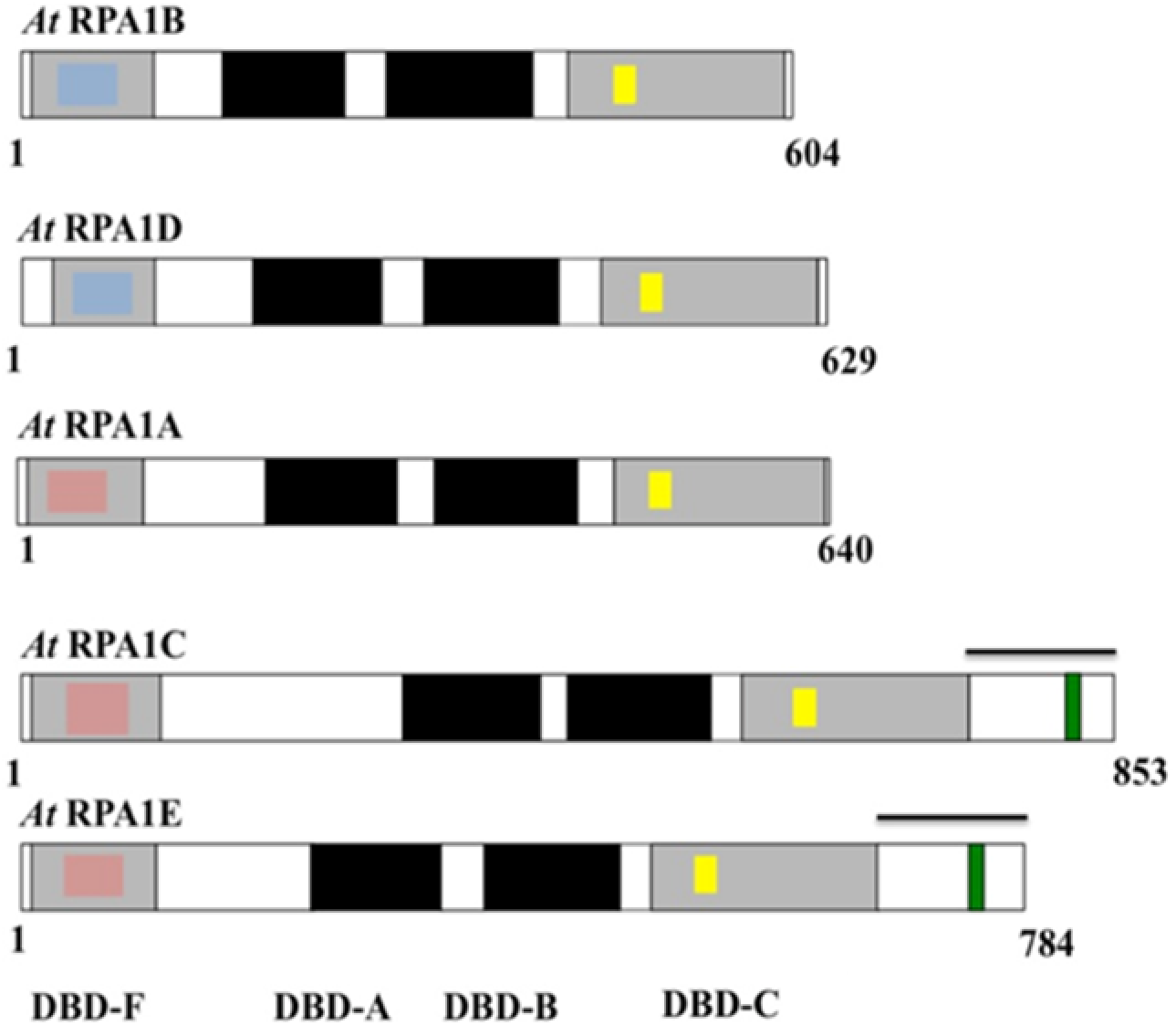
Model of Arabidopsis thaliana RPA1 paralogs. *Arabidopsis thaliana* has five RPA1 paralogs. The paralogs all share the same four DNA binding domains (DBD-A, DBD-B, DBD-C, and DBD-F) but Binding Surface I (BS-I, pink boxes) is only found in RPA1A, RPA1C, and RPA1E. DBD-F in RPA1B and RPA1D does not contain the BS-I subdomain (blue boxes). All RPA1 paralogs contain a zinc finger motif (ZFM) within DBD-C (yellow boxes). RPA1C and RPA1E also have a unique C-terminal extension which contains a CCHC-type type ZFM (green; Aklilu & Culligan, 2016)

In animals and yeast, the DNA damage response involves the hyper-phosphorylation of RPA, but this is not the case in plants (Binz et al., 2004; Vassin et al., 2004). Indeed, following induced DNA damage in rice, there was no hyper-phosphorylation detected (Marwedel et al., 2003). Given the lack of hyper-phosphorylation, another method of regulation is needed and an interaction with the RPA1 C-terminal extension ZFM is a prime candidate for plant-based RPA regulation, as it is unique to plants and only present in paralogs involved in DNA damage repair.

To determine the role of the C-terminal ZFM of RPA1C and RPA1E, we sought to specifically remove each respective ZFM from RPA1C and RPA1E in WT (Col-0) Arabidopsis. To achieve this, a CRISPR-Cas9 gene editing system was used to generate Arabidopsis plants that had the C-terminal extension ZFM removed from either RPA1C or RPA1E (Hereafter referred to as ZFKO lines). To observe the impact of the removal of the ZFM on the function of RPA in DNA damage repair, the ZFKO lines were challenged with DNA damaging agents, and their susceptibility was compared to both WT and to previously characterized T-DNA null mutants of RPA1C and RPA1E (*rpa1c* and *rpa1e*). To further characterize the ZFKO lines they were also crossed with GUS lines which are activated specifically by either SSA or SDSA. Using these generated GUS lines, the capability of the ZFKO lines to function in specific DSB repair pathways could be assessed and the exact role(s) of the ZFMs could be more precisely determined. In this study we find that the ZFKO lines behaved identically to their respective null lines both when challenged with DNA damaging agents and when tested for activity in HRR pathways. These results indicate that the ZFM is crucial to the functionality of the RPA1C and RPA1E paralogs in DNA damage repair.

## Materials and Methods

### Plant Materials and Growth

Salk T-DNA insertion mutants (Salk IDs: rpa1c, Salk_085556; rpa1e, Salk_120368) were obtained from the *A. thaliana* Biological Resource Center (ABRC). Homozygous mutants for these lines were identified via PCR using gene and T-DNA specific primers. Ecotype Columbia (Col-0) was employed as the WT control. Prior to germination, seeds were surface sterilized by agitation in a solution of 20% bleach and 0.02% Tween-20 for five minutes. Seeds were then rinsed with autoclaved double-distilled water three times, with further agitation used during each rinse. Following sterilization, seeds were sown on nutrient phytoagar plates containing 1x MS salts (PlantMedia, Dublin, Ohio, USA) pH 5.7, 0.05 g/L MES and 1.0% (w/v) phytoagar (PlantMedia, Dublin, Ohio, USA). Seeds were stratified at 4°C for two days in the dark before plates were placed vertically in a growth chamber under cool-white lights at an intensity of 100-150 mmol/m^2^/sec at 22°C and a photoperiod of 16hr light/ 8hr dark. After approximately one week of growth, the plants were transferred to soil growth medium (SUNGRO Horticulture, Seba Beach, Canada) with each plant given its own pot. Plants were watered every 3 days with tap water supplemented by Miracle-Gro® 15-30-15 plant fertilizer at a concentration of 0.45 g/liter (Scotts Miracle-Gro products Inc., Marysville Ohio, USA).

### DNA Damage Hypersensitivity Assays

Approximately 50 surface-sterilized WT or mutant seeds were sown on 3 replicate plates containing 1X MS phytoagar media with or without camptothecin (CPT) (SIGMA, St. Louis, MO, USA) at a concentration of 15nM. Seeds were stratified and then grown as previously described. Plates were photographed, and root length measured following 11 days of growth. Root measurements were of primary root growth from the root junction to the root tip.

For gamma-radiation assays, Arabidopsis seeds or plants were irradiated using a Cs 137 source (Massachusetts Institute of Technology, Cambridge, MA, USA), dose rate 81 rad/minute. Sterilized seeds were imbibed in water at 4°C for 2 days, irradiated to a dosage of 200 Gy (20,000 rad) and then immediately placed on 1X MS phytoagar plates for germination in the growth chamber. For seedlings, plate-grown 5-day-old seedlings were irradiated to a dosage of 100 Gy and then immediately returned to the growth chamber. For both the experiments using seeds and seedlings, growth continued until an age of 11 days, at which point the plants were photographed and primary root growth was measured. To calculate the relative percent growth reduction of the treated plants compared to the untreated plants we used the formula:

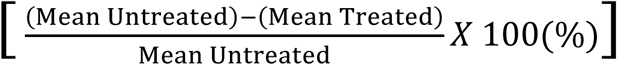

### Gus Assay

Plant lines carrying IU.GUS, DGU.US and I-SceI were acquired courtesy of H. Puchta (Mannuss et al., 2010). Lines were confirmed as homozygous via gene-specific primers. All three acquired lines were crossed with the following Arabidopsis lines: *rpa1c*, *rpa1e*, C-ZFKO1, E-ZFKO and the F1 offspring were allowed to self-fertilize to produce F2 plants homozygous for a mutant RPA1C or RPA1E background and either I-SceI or a GUS reporter. Finally, crosses between F2 lines yielded heterozygous plants for both I-SceI and one GUS reporter line in a homozygous RPA1C or RPA1E mutant background.

Prior to application of the X-Gluc reagent, plants were fixed in cold 90% acetone for 20 minutes. Following fixation, plants were incubated in 50 mM NaPO_4_, pH 7.2, 0.5 mM K_3_Fe(CN)_6_, 0.5 mM K_4_Fe(CN)_6_, 10 mM EDTA, 0.01% Trition X-100, and 2 mM X-Gluc (Gold Biotechnology, St. Louis, MO, USA) overnight at 37°C. Plants were decolorized in 95% ethanol for 3×30 minutes. Plants were directly imaged under an Olympus SZ51 light microscope. Quantitative evaluation of GUS staining was completed by examining leaves of at least four plants per genotype per trial. Each individual instance of coloration was counted as a single “speck” (Mannuss et al., 2010). Results were averaged to find the number of specks per leaf for each line. Values were compared via one-way ANOVA followed by LSD to determine significant differences between groups.

### CRISPR/Cas9 Mediated Genome Editing

Plasmids for CRISPR/Cas9 genome editing were constructed from the vectors pEG_302v2, pEN_Comaira.1, pEN_Comaira.2, and pEN-RC9.3 using Gateway Cloning. Successful cloning was confirmed via sequencing prior to transformation of plasmids into electrocompetent *Agrobacterium tumefaciens* GV3101. Arabidopsis plants were transformed using the floral dip technique and resulting seeds were screened using growth media containing 10 mg/L Glufosinate ammonium (Clough & Bent, 1998). Further confirmation of successful genome editing was done with PCR with gene specific primers and DNA sequencing.

## Results

### C-terminal deletions of the Zinc-Finger Motif (ZFM) in RPA1C and RPA1E were constructed using CRISPR-Cas9

To determine the role of the ZFM of RPA1C and RPA1E, a CRISPR/Cas9 system was employed to specifically remove these regions from WT (Col-0) Arabidopsis RPA subunit genes. 8 total crRNAs were designed and inserted into plasmids and further transformed into *Agrobacterium tumefaciens* GV3101. The Agrobacterium was used for floral dip plant transformations and the resulting T1 seeds were screened for resistance to the herbicide Glufosinate ammonium. Resistant seeds were screened with PCR to identify plants with any modification to the C-terminal region of the RPA subunit. Plants with altered C-terminal regions were then sequenced to confirm the successful genetic editing. Three positive lines were used for the subsequent trials, two RPA1C mutant lines (referred to as C-ZFKO1 and C-ZFKO2) and one RPA1E mutant line (referred to as E-ZFKO) (Table 1).

**Table 1.** Zinc finger deletions used in this study.

| crRNA | Expected Deletion Size (bp) | Actual Deletion Size (bp) |
| --- | --- | --- |
| C-ZFKO1 | 206 | 202 |
| C-ZFKO2 | 160 | 225 |
| E-ZFKO | 160 | 159 |

### Arabidopsis RPA1 Zinc-finger deletion lines (ZFKO) are hypersensitive to camptothecin-induced DNA damage

To test the ZFKO lines for functionality during DNA double-strand break repair, we employed seedling hypersensitivity assays in response to the topoisomerase inhibitor camptothecin (CPT). CPT induces DSBs by inhibiting the re-ligation of DNA topoisomerase I during replication (L. F. Liu et al., 2000). We previously tested CPT on RPA1 T-DNA null mutants, finding that *rpa1c* lines were hypersensitive (Aklilu et al., 2014). As shown in Figure 3A, in the absence of camptothecin all lines exhibited identical growth phenotypes. However, upon exposure to camptothecin all null mutant and ZFKO lines displayed significantly reduced growth (Figure 3B). RPA1C null and ZFKO mutants were particularly sensitive and had ∼67% reduced root length compared to WT (Figure 4). RPA1E null and ZFKO mutants displayed a lesser, but still significant, degree of sensitivity with ∼25% reduction of root length versus WT. In all cases, the ZFKO lines and their respective null mutants displayed identical phenotypes, indicating that the ZFM of both RPA1C and RPA1E is crucial to the overall functionality during DNA damage repair. Furthermore, the increased sensitivity seen in RPA1C mutant lines suggests that RPA1C plays a more substantial role in DNA damage repair than RPA1E.

**Figure 3.**
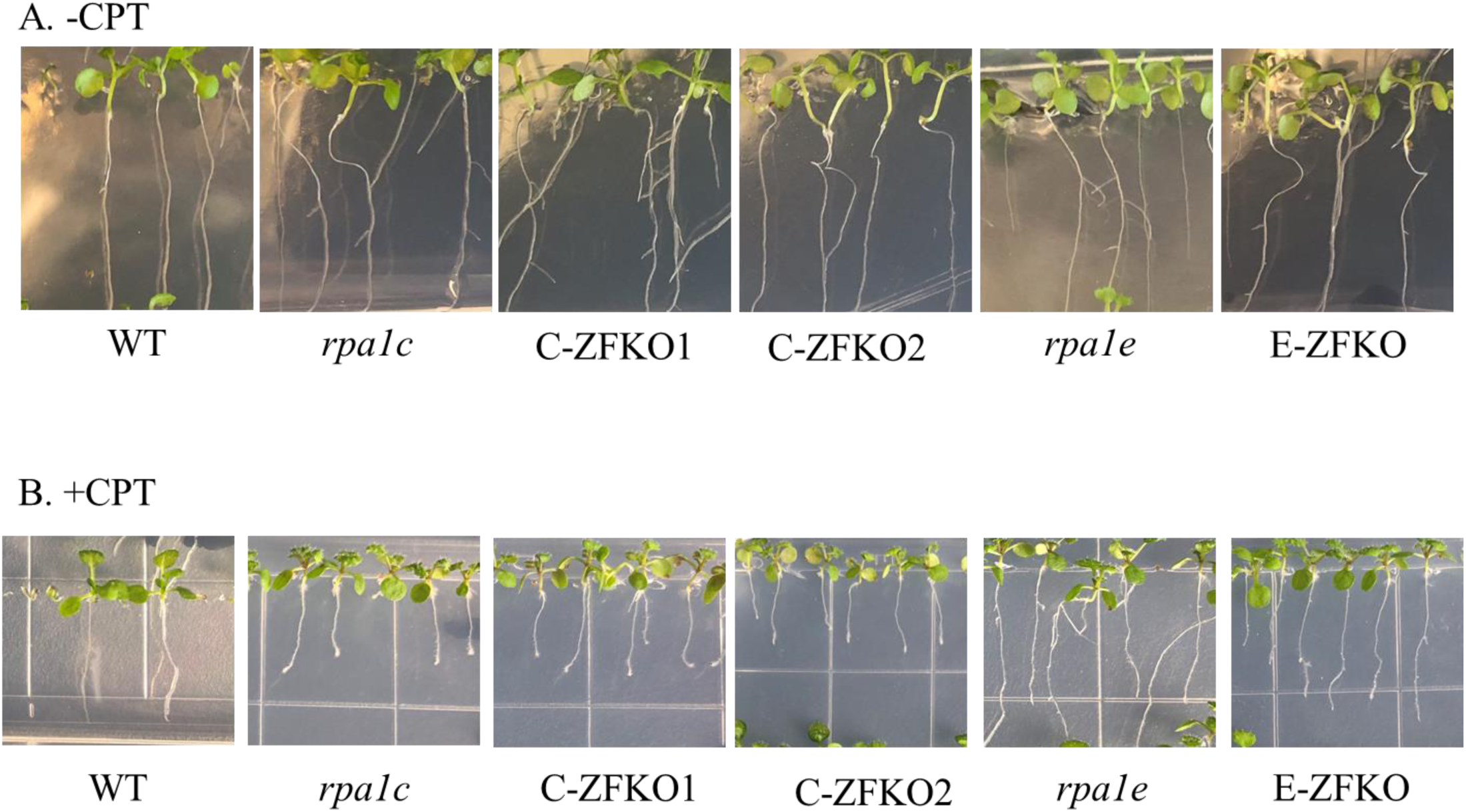
Hypersensitivity assay of RPA1 mutants to damage caused by CPT. 11-day-old WT and *rpa1* null and ZFKO mutant seedlings grown on MS medium supplemented with either 0 nM (A) or 15 nM CPT (B).

**Figure 4.**
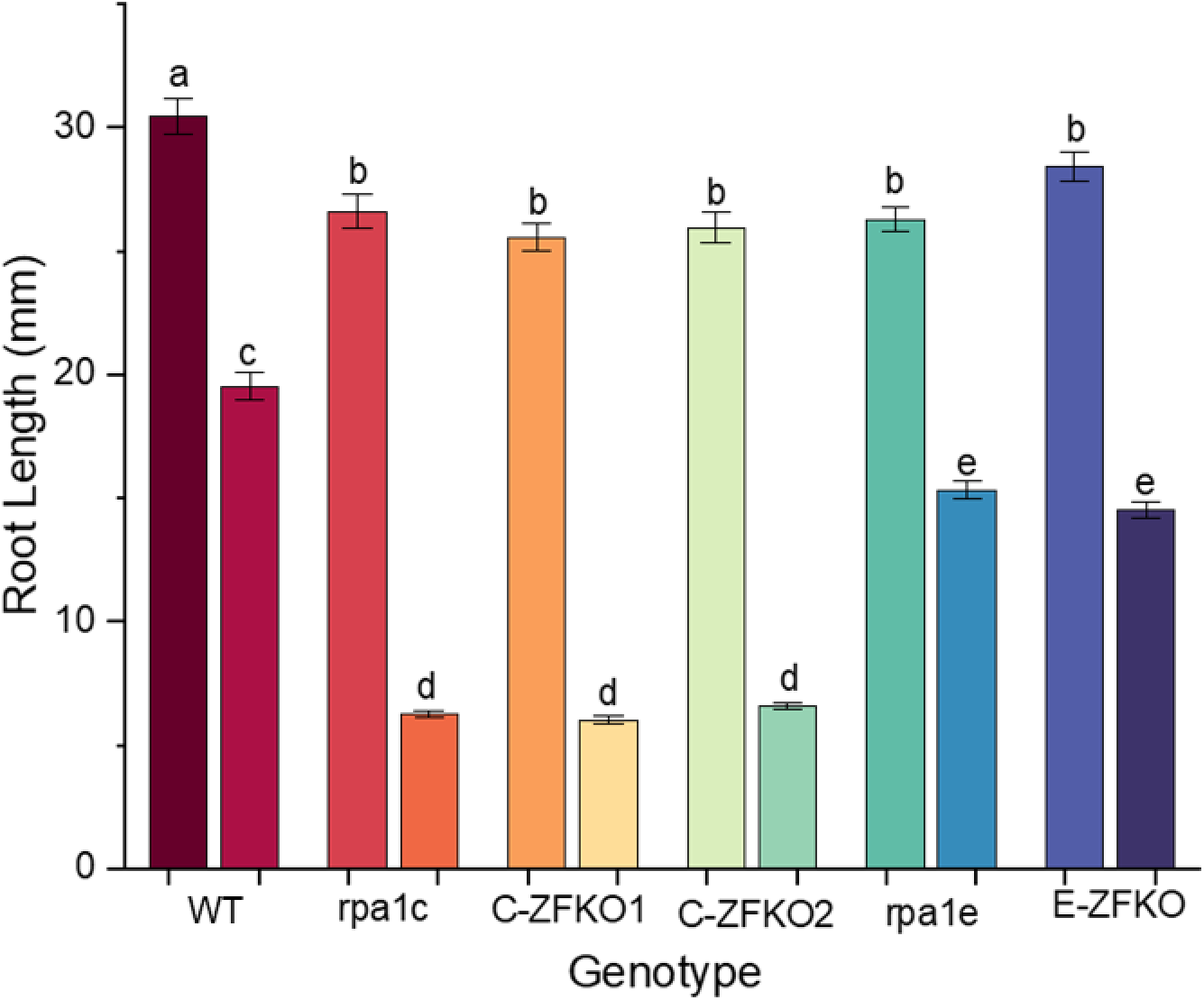
Root length measurements of all control (left) and experimental (right) plants used in CPT exposure experiment. Statistical analysis employed F-test (ANOVA) and LSD at P ≤ 0.05. Error bars denote standard error and bars with different letters indicate significant differences (n>60).

### ZFKO lines are hypersensitive to gamma-induced DNA double-strand breaks

DNA damage caused by CPT is limited both in terms of the type of damage (double-strand breaks) and the stage of the cell cycle (S/G2 phase) in which it acts (L. F. Liu et al., 2000). To examine the functionality of the group C RPA1 paralogs in response to more generalized DNA damage, we employed gamma radiation as an additional source of DNA double-strand breaks. Gamma radiation induces DSBs as well as SSBs and abasic sites and is not restricted to S/G2 phase. Plants were exposed to 100 Gy of gamma radiation as 5-day-old seedlings or exposed to 200 Gy of gamma radiation prior to germination. In both cases, root length measurements were assessed after 11 days of growth.

Under normal growth conditions there were no observed differences in root length for any of the genotypes (Figures 5 and 7). However, RPA1C null and ZFKO mutant seedlings exposed to 100 Gy of gamma radiation showed a ∼18% decrease in root length compared to WT (Figure 6). RPA1E null and ZFKO mutant seedlings were not significantly different than WT. Further experiments were conducted using a higher dose of radiation applied prior to germination and produced similar results. RPA1C null and ZFKO mutant seeds exposed to 200 Gy of gamma radiation showed a ∼15% decrease in root length compared to WT while RPA1E null and ZFKO mutant seedlings were not significantly different than WT (Figure 8). These data indicate that RPA1C is crucial to all types of DNA damage repair while RPA1E may have a more specialized role. Additionally, the null and ZFKO lines for both RPA1C and RPA1E continued to exhibit the same phenotype in response to DNA damage, indicating that the C-terminal extension zinc finger is necessary for all types of DNA damage repair activity.

**Figure 5.**
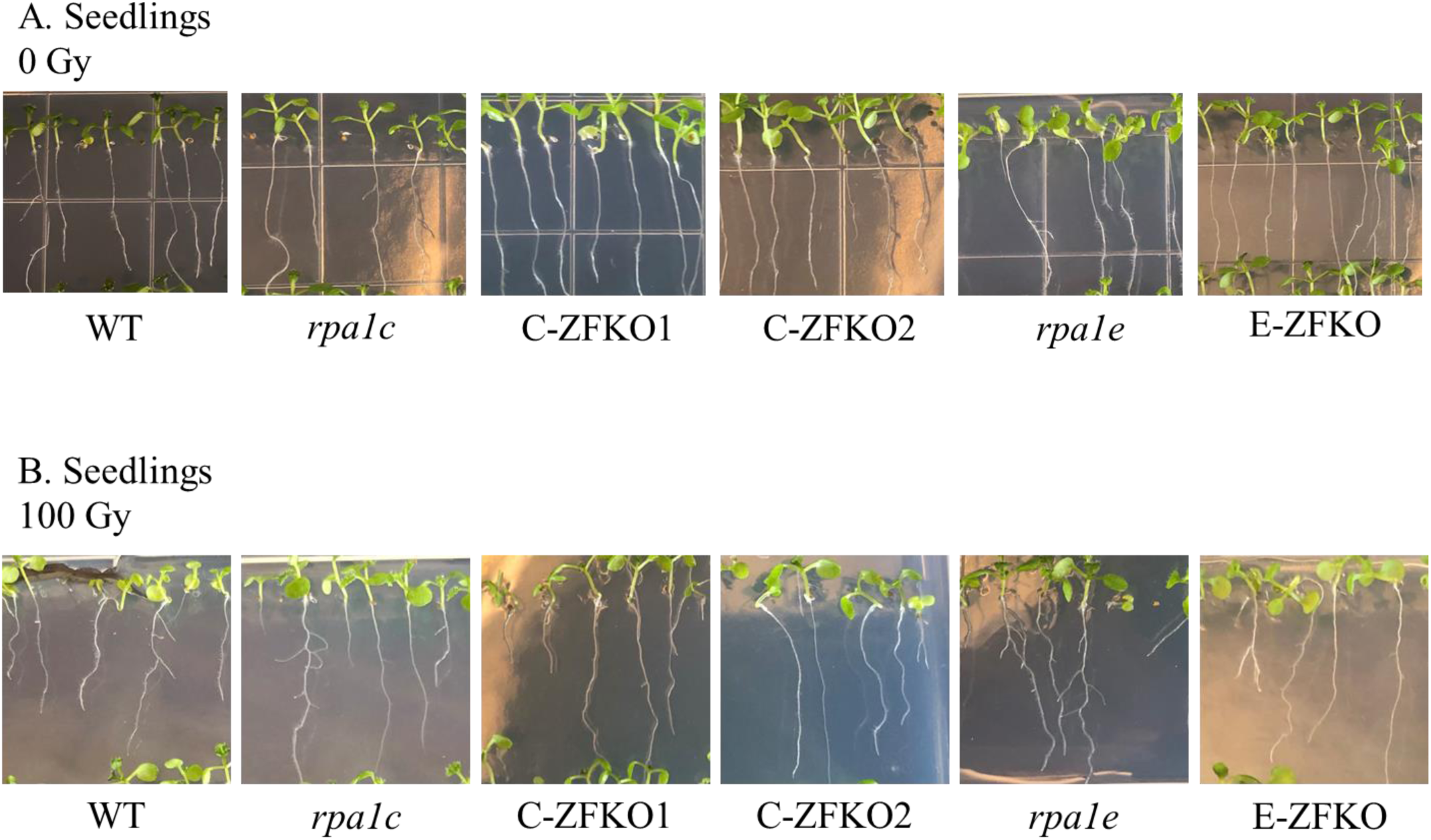
Hypersensitivity assay of RPA1 mutant seedlings to damage caused by gamma radiation. 11-day-old WT and *rpa1* null and ZFKO mutant seedlings grown on MS medium and treated with 0 (A) or 100 (B) Gy of gamma radiation when 5 days old.

**Figure 6.**
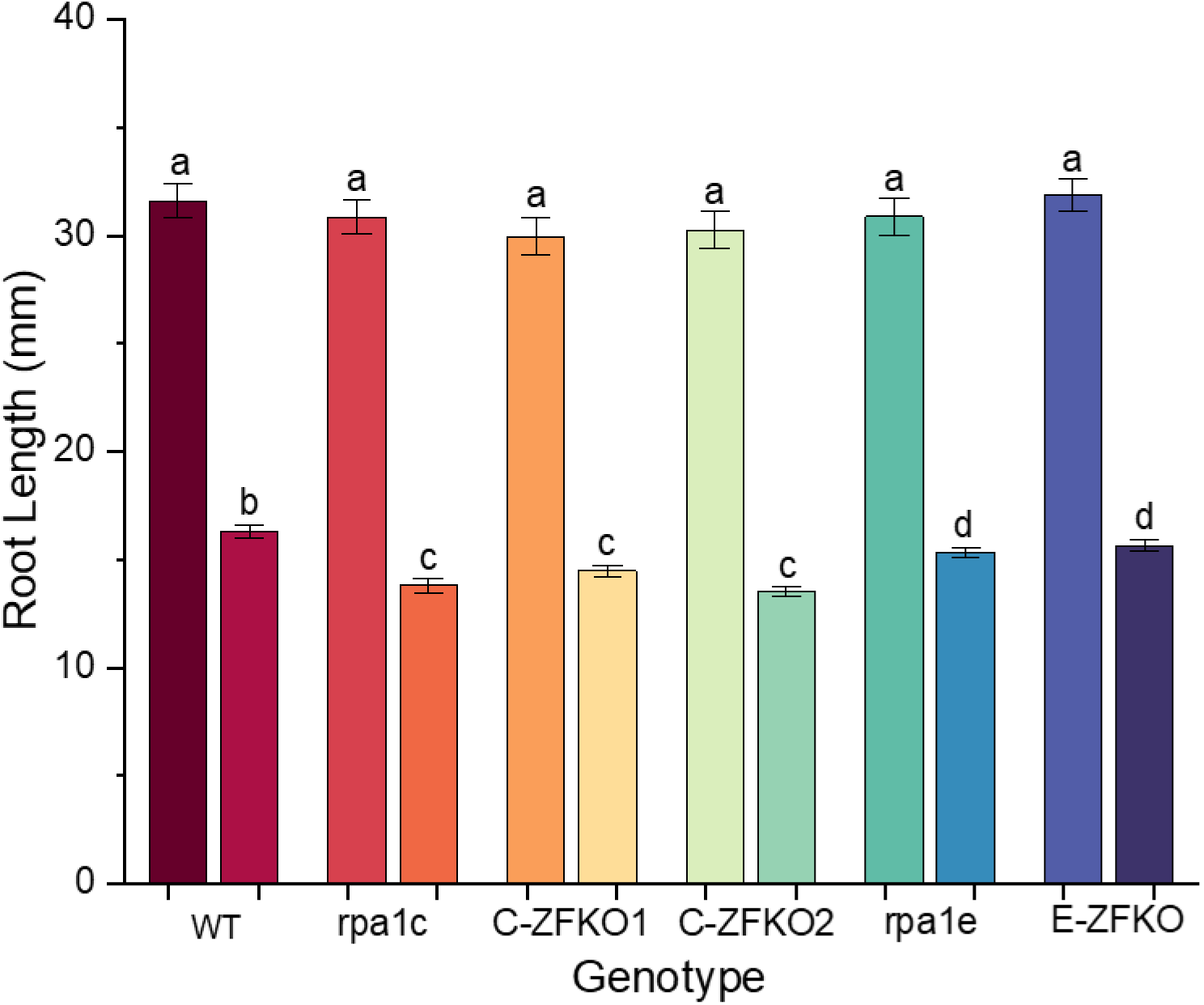
Root length measurements of all control (left) and experimental (right) plants exposed to gamma radiation as seedlings. Statistical analysis employed F-test (ANOVA) and LSD at P ≤ 0.05. Error bars denote standard error and bars with different letters indicate significant differences (n>60).

**Figure 7.**
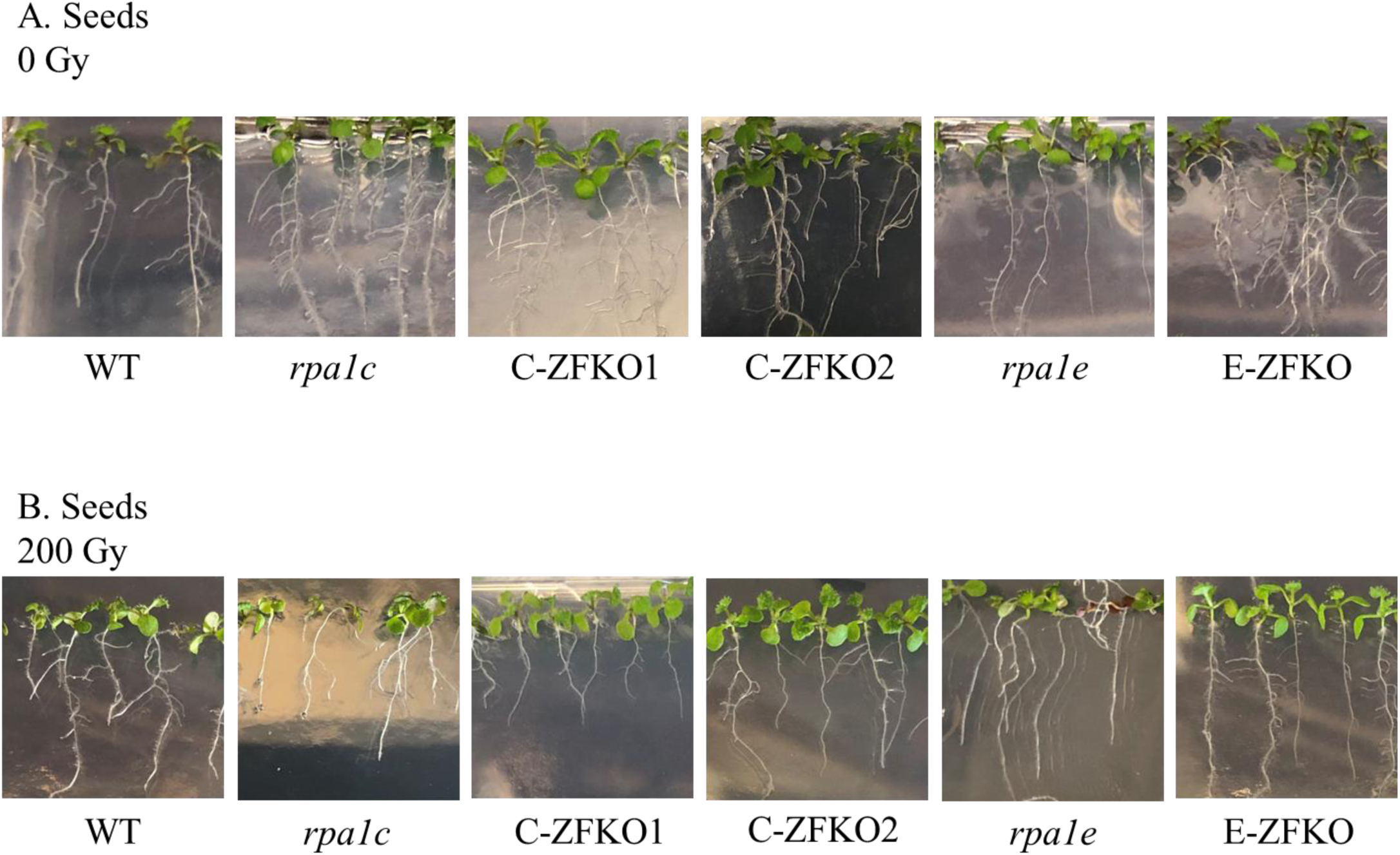
Hypersensitivity assay of RPA1 mutant seed to damage caused by gamma radiation. 11-day-old WT (Col-0) and *rpa1* null and ZFKO mutant seedlings grown on MS medium and treated with 0 (A) or 200 (B) Gy of gamma radiation as seeds immediately prior to sowing and transfer to the growth chamber.

**Figure 8.**
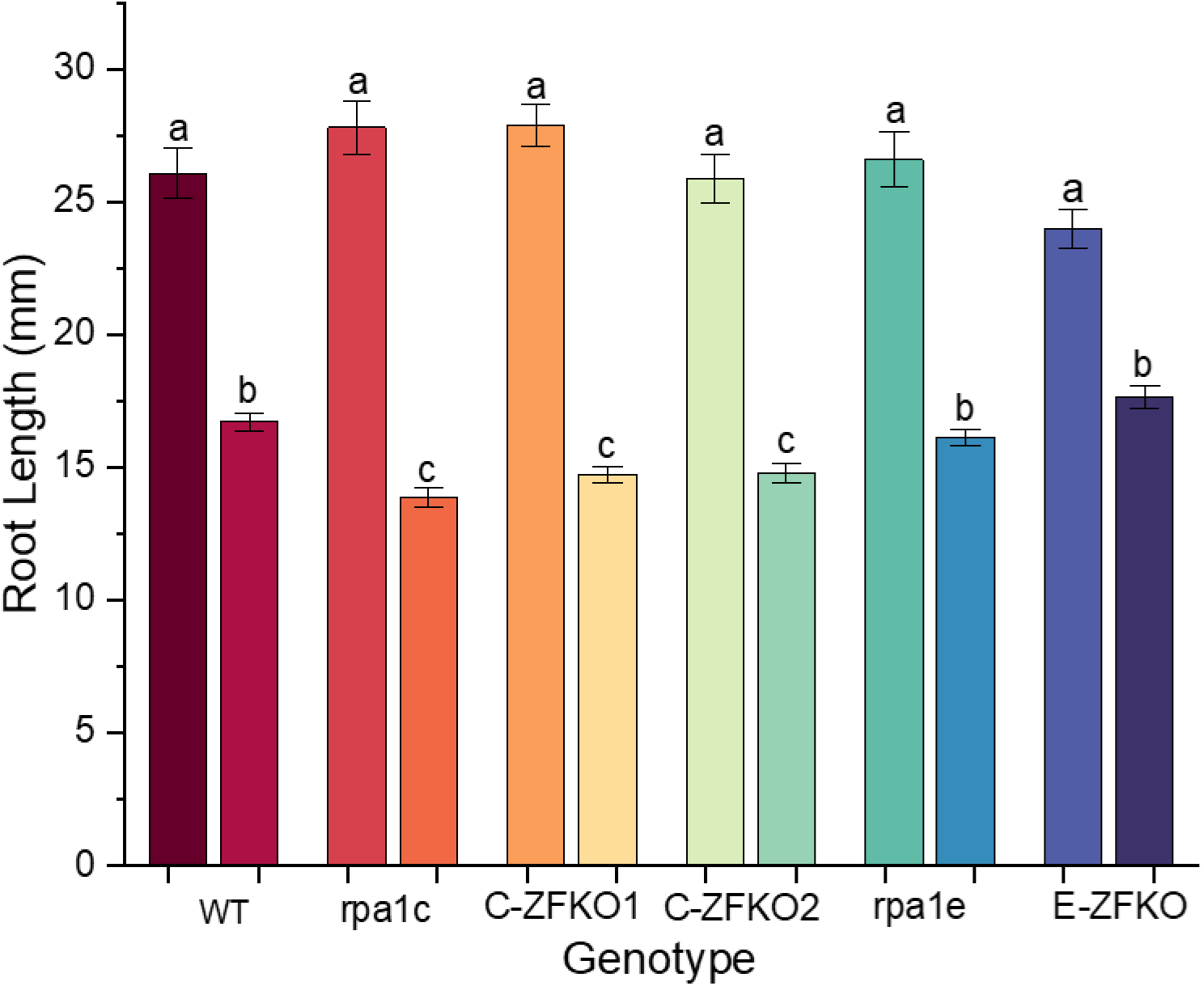
Root length measurements of all control (left) and experimental (right) plants exposed to gamma radiation as seeds. Statistical analysis employed F-test (ANOVA) and LSD at P ≤ 0.05. Error bars denote standard error and bars with different letters indicate significant differences (n>60)

### Both C-ZFKO and E-ZFKO lines have greatly reduced homologous recombination repair activity

Studies employing DNA damaging agents (as above) demonstrate that the C-terminal ZFM is crucial for the function of RPA during DNA repair. However, it is unclear whether the paralogs may promote or inhibit specific HRR pathways. It is also possible that the absence of the C-terminal ZFM would result in variations in HRR distinct from that seen in either RPA1 null mutants or WT. To address this, RPA1 null lines and ZFKO lines were crossed with GUS reporter constructs DGU.US and IU.GUS to measure HRR pathway activity by RPA1 paralogs during DSB repair (Orel et al., 2003).

The resulting lines were either RPA1 null mutants or ZFKO mutants combined with an open-reading frame for the restriction enzyme I-SceI and either the DGU.US or IU.GUS GUS reporter constructs (Figure 9) (Mannuss et al., 2010). The GUS constructs are activated by I-SceI restriction and further repair by either SSA (DGU.US lines) or SDSA (IU.GUS lines) (Orel et al., 2003). Once repair occurs, GUS expression results in the production of blue pigment specks (through the hydrolysis of X-Gluc substrate). The relative level of pigment produced in each line was quantified by counting unique repair events (blue specks) per leaf then calculating the average number of events per leaf for each genotype to determine the level of HRR activity in that line.

**Figure 9.**
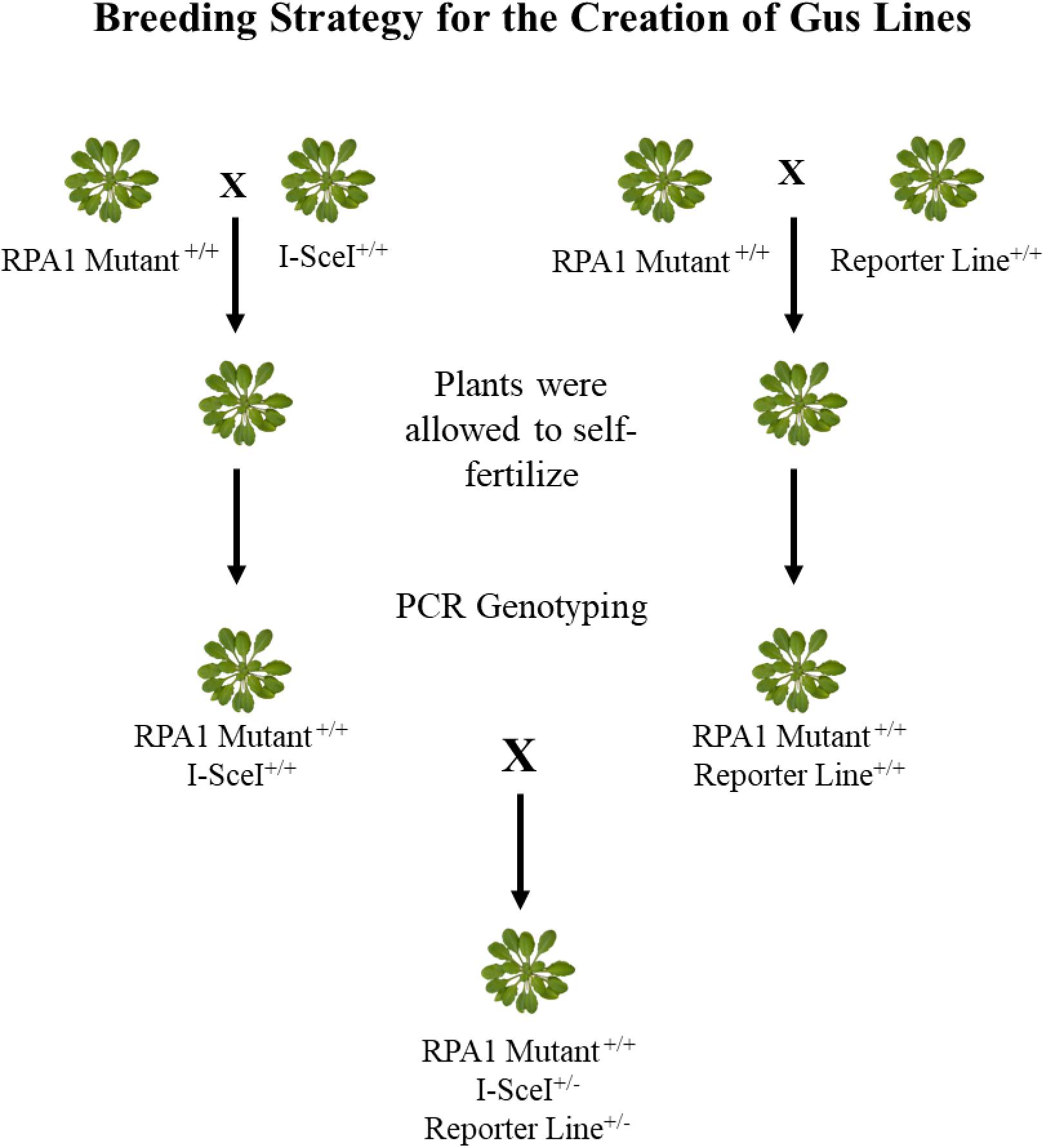
Model of the breeding strategy used to generate GUS lines with mutant RPA1 backgrounds.

Compared to WT, all mutant lines showed a reduction in SSA activity (Figures 10 and 11). The reduced activity was more prominent in the RPA1C null and C-ZFKO lines, which both showed a ten-fold reduction in SSA activity. The RPA1E null and E-ZFKO lines had a less drastic response but still showed an approximate two-fold reduction. This indicates that both RPA1C and RPA1E are involved in SSA-specific DSB repair, with RPA1C playing a leading role over RPA1E.

**Figure 10.**
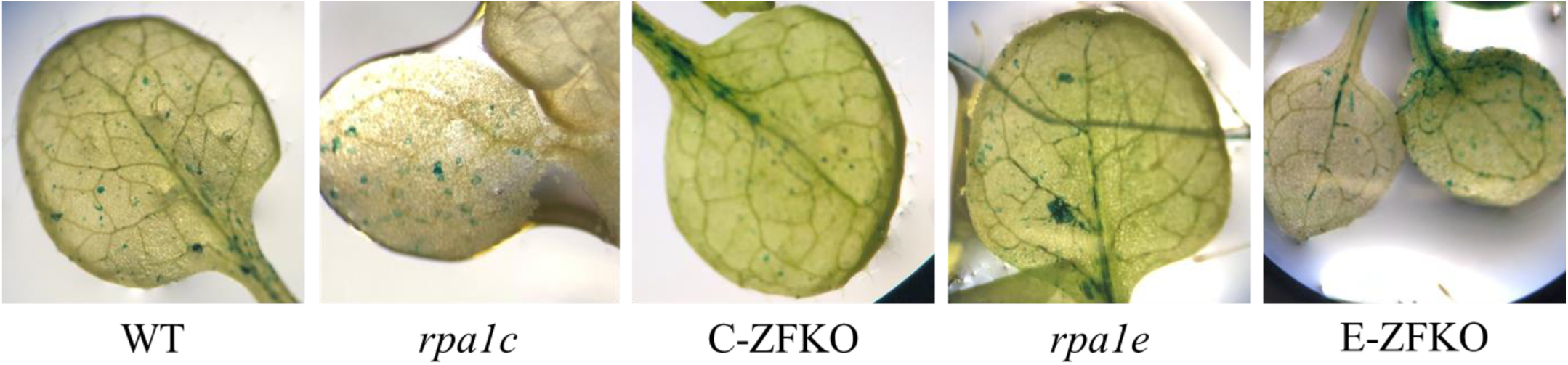
Representative images of GUS activity in single-strand annealing GUS assay. The level of activity was quantified by counting and averaging the number of individual specks per leaf.

**Figure 11.**
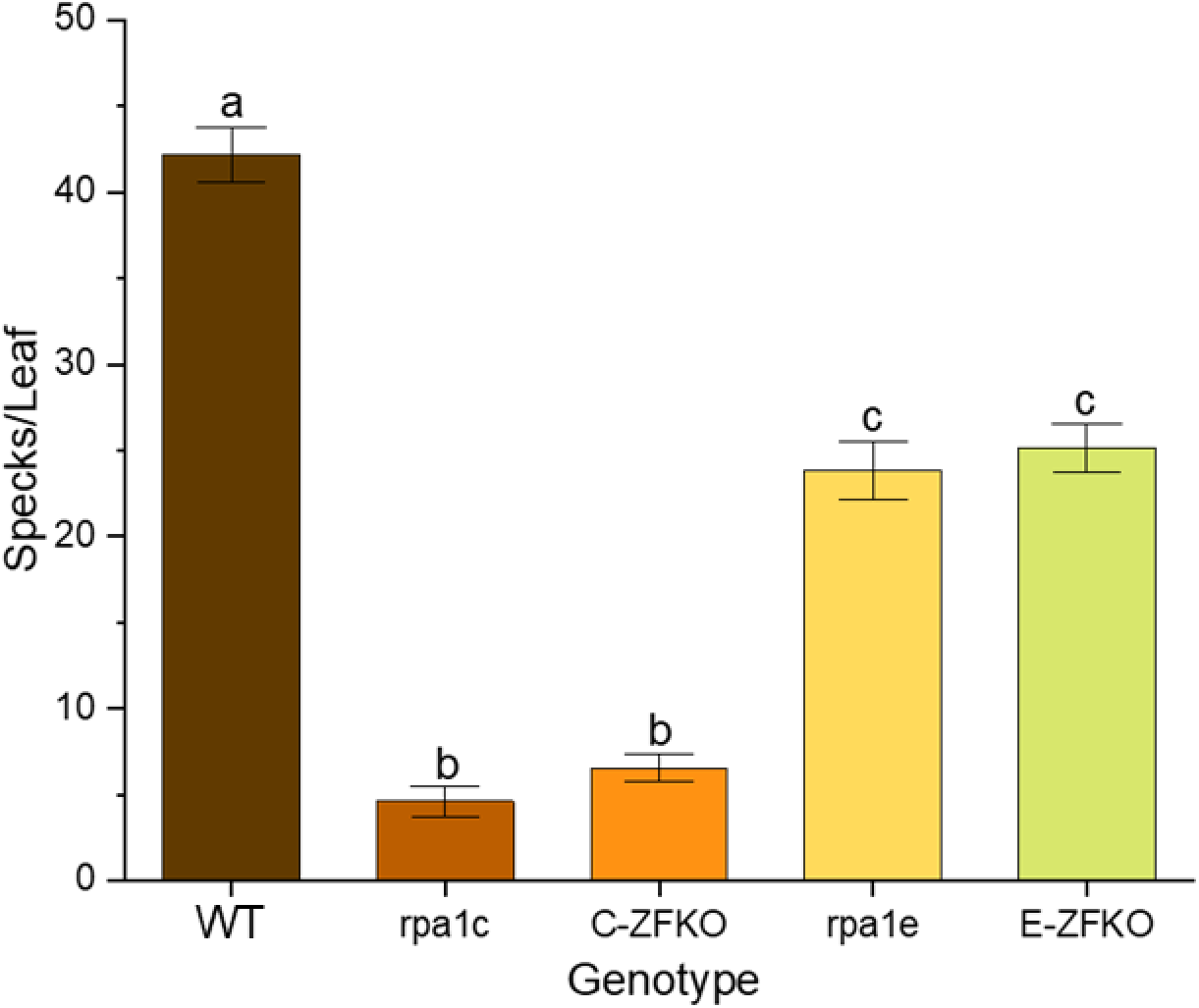
Single-strand annealing (SSA) activity of two-week-old seedlings. The level of activity was quantified by counting and averaging the number of individual specks per leaf. Statistical analysis employed F-test (ANOVA) and LSD at P ≤ 0.05. Error bars denote standard error and bars with different letters indicate significant differences (n>40).

In contrast, RPA1C and RPA1E null lines and ZFKO lines all display a similar reduction in SDSA activity compared with WT (Figures 12 and 13). All four lines show an approximate two-fold reduction in SDSA compared to WT, suggesting that both RPA1C and RPA1E function similarly in SDSA repair pathways.

**Figure 12.**
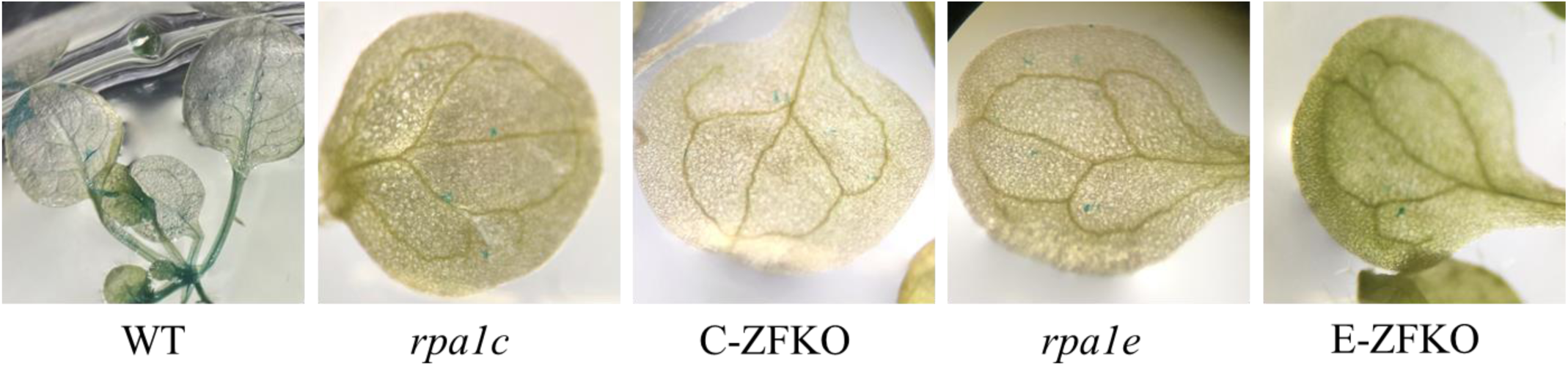
Representative images of GUS activity in synthesis-dependent strand annealing GUS assay. The level of activity was quantified by counting and averaging the number of individual specks per leaf.

**Figure 13.**
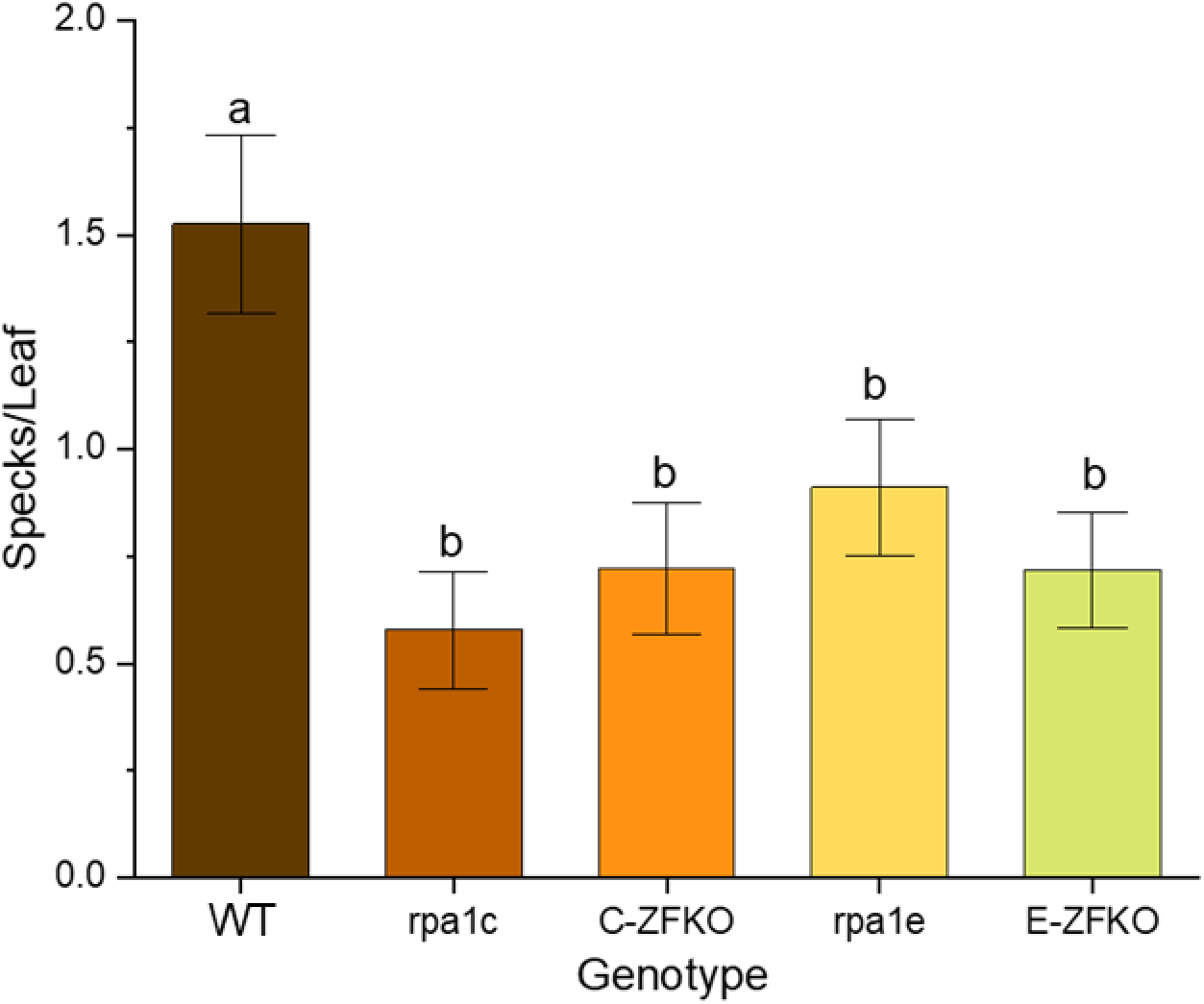
Synthesis-dependent strand (SDSA) activity of two-week-old seedlings. The level of activity was quantified by counting and averaging the number of individual specks per leaf. Statistical analysis employed F-test (ANOVA) and LSD at *P* ≤ 0.05. Error bars denote standard error and bars with different letters indicate significant differences (n>40).

Viewing the combined results of the GUS assays, the RPA1C null and C-ZFKO lines showed a drastic reduction in SSA activity (ten-fold reduction) as well as a modest decrease in SDSA activity (two-fold reduction). In contrast, the RPA1E null and E-ZFKO lines showed a modest reduction (two-fold) for both SSA and SDSA (Figure 14). The greater reduction of SSA compared to SDSA seen in the RPA1C and C-ZFKO lines indicates a possible preference for SSA following binding of RPA1C during DSB repair, whereas RPA1E binding does not seem to affect the choice of HRR pathway. Additionally, in each experiment, the RPA1C null line and C-ZFKO behaved identically, as did the RPA1E null line and E-ZFKO. This indicates that the C-terminal extension zinc finger motif is crucial to the functionality of both RPA1C and RPA1E in DSB repair using both SSA and SDSA and its removal does not lead to the favoring of either repair pathway.

**Figure 14.**
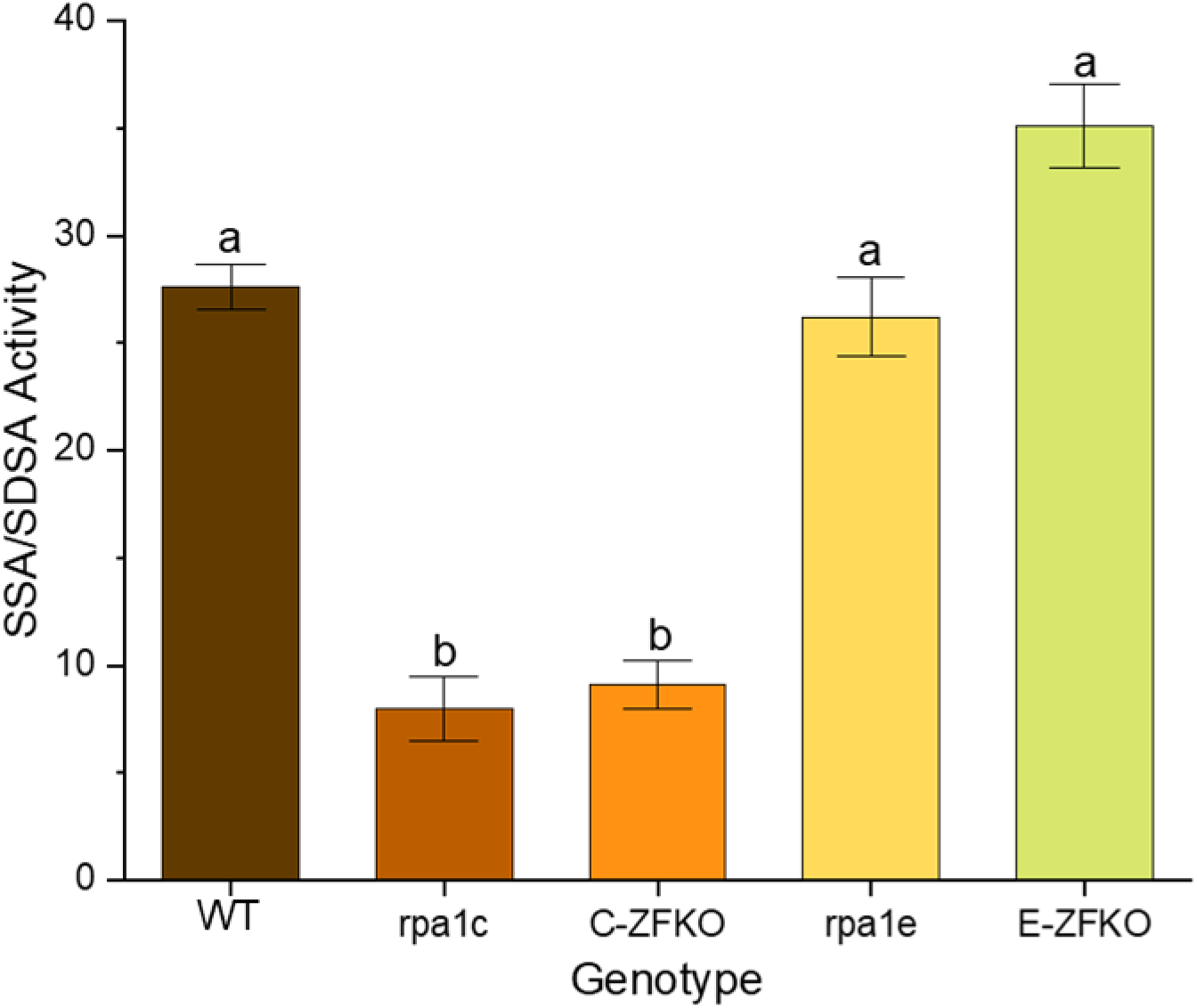
Relative levels of single-strand annealing (SSA) and synthesis-dependent strand annealing (SDSA) activity. Statistical analysis employed F-test (ANOVA) and LSD at *P* ≤ 0.005. Error bars denote standard error and bars with different letters indicate significant differences (n>40).

## Discussion

In this study, we find that the C-terminal extension of the group C paralogs and specifically the CCHC-type ZFM within the C-terminal extension is crucial to the functionality of the paralogs during DNA damage repair. The removal of the ZFM resulted in hypersensitivity identical to the phenotypes of RPA1 null mutants (*rpa1c* and *rpa1e)* in response to damage from both CPT and gamma radiation. Furthermore, the RPA1C null and ZFKO lines both displayed significantly greater susceptibility to DNA damage upon exposure to both CPT and gamma radiation treatment than the RPA1E null and ZFKO lines. These results are consistent with previous studies showing RPA1C plays a more significant role over RPA1E in DNA damage and repair responses (Aklilu & Culligan, 2016). This supports a hypothesis in which RPA1E has a more specialized role in DNA damage repair compared to a more general role of RPA1C (Aklilu et al., 2014). One possible scenario is that RPA1E functions preferentially during certain phases of the cell cycle or in response to a specific type of DNA damage.

We further investigated the specific roles of the group C RPA1 paralogs employing a GUS system to measure levels of SSA and SDSA in RPA1C and RPA1E mutant lines. When observing specific DSB repair pathways the *RPA1C* null line (*rpa1c*) displayed a larger decrease in SSA activity versus the *RPA1E* null line (*rpa1e*). However, no difference was observed between mutant lines when observing SDSA activity. Furthermore, CRISPR-generated Arabidopsis lines lacking the C-terminal extension ZFM from RPA1C or RPA1E displayed identical phenotypes to their respective null mutant. These data indicate that both RPA1C and RPA1E are involved in both HRR repair processes. The greater decrease in SSA activity in *RPA1C* mutants compared to *RPA1E* mutants could indicate a preference towards SSA following RPA1C binding but could also reflect the overall greater importance of RPA1C in DSB repair. It should be noted that this result differs somewhat from previously published work, which did not state any significant difference in the rate SSA between RPA1C and RPA1E null mutants (Liu et al., 2017). However, the Liu study was not focused on the variation between RPA1 paralogs, so no direct comparative assay was conducted. Additionally, the Liu study did not investigate the rates of SDSA in any of the Arabidopsis lines studied so it is unknown whether those results would align. Regardless, the identical phenotypes of the RPA1 C null/C-ZFKO lines and the RPA1E null/E-ZFKO lines in our study also make our findings more reliable as it is unlikely both lines would display the same phenotype by chance. These data, combined with the data from the DNA damage sensitivity assay, supports RPA1C having a larger role than RPA1E in DNA damage repair. As with the DNA damage assays, the phenotype of the null mutants was identical to the corresponding ZFKO lines, indicating the functionality of both RPA1C and RPA1 in both HRR pathways is reliant upon the presence of the C-terminal extension ZFM.

While the work presented here does show the importance of the C-terminal extension ZFMs, the exact role of the ZFMs is still unclear. CCHC-type ZFMs have been shown to have particularly high binding affinity to single-stranded nucleic acids and particularly ssRNA, meaning that a regulatory role might be possible (Wang et al., 2021). However, a regulatory role alone may not adequately explain the observed phenotype, which was equivalent to a null mutant. Another possibility is that the ZFM works in conjunction with BS-I and aids in binding to DNA during DNA damage repair. It could accomplish this by interacting directly with the DNA, but CCHC ZFMs have also been noted as being capable of binding to proteins (Liew et al., 2000; Matthews et al., 2000; Tapia-Ramírez et al., 1997). Therefore, it is possible that the ZFMs are involved in the recruitment of other proteins such as Rad51 and Rad54 to the site of DNA damage. Additionally, as the C-terminal extension is unique to plants it is possible that the binding target is a plant specific factor and that the ZFM and its binding capabilities are related to the differential regulation of RPA between plant and other eukaryotes. Regardless of the exact function(s) of the ZFMs it is clear that they are crucial to the functionality of the group C RPA1 subunits and merit further investigation.

